# Genetic hubs: a phylogeographer’s widget for pinpointing of ancestral populations

**DOI:** 10.1101/419796

**Authors:** Mikula Ondřej

## Abstract

Statistical phylogeography benefits from the development of increasingly realistic models of spatially structured genetic variation. Their fitting, however, is computationally demanding and requires population and/or genomic sampling that is not available for many species of interest. ‘Genetic hubs’ is a method that can be used for exploratory analyses of various kinds of genetic data, including those as typical in mitochondrial phylogeography, i.e. many small samples of single locus genotypes scattered throughout the species distributional range. ‘Genetic hubs’ allows to quantify and visualize gradients of genetic variation with the aim to pinpoint possible origin of expansion. It estimates local genetic variability as an accessibility of all genetic variation from the site in question and it allows to take dissimilarity of genotypes into account. The method represents fast and versatile tool that can be used whenever history of range expansion is assumed to shape the observed distribution of genetic variation and it is useful especially for preliminary analyses whose purpose is to provide sound basis for formulation of testable hypotheses and design of follow-up studies.

## Background

Phylogeography attempts to understand processes and historical events that shaped spatial structure of genetic variation and, compared to other branches of population genetics, it focuses on their timing and geographical setting (Avise, 2000; Knowles & Maddison, 2002). One important goal in such endeavor is to estimate the origin of expansion, i.e. the location of ancestral population, from which the rest of distribution range was colonized.

The range expansion results in a serial founder effect that can be thought as a spatial analog of genetic drift (Slatkin & Excoffier, 2012), which creates gradients of decreasing genetic diversity, increasing linkage disequilibrium and flattening ancestral allele frequency spectrum. All these phenomena can be used when searching for the origin of expansion of humans and the observed trends match expectations based on paleontologically well-evidenced out-of-Africa dispersal scenario (DeGiorgio, Jakobsson & Rosenberg, 2009; Jakobsson et al., 2008; Ramachandran et al., 2005). These analyses could be performed because of at least hundreds of loci genotyped in hundreds of individuals from tens of populations. The source population can be also identified by analysis of fewer loci genotyped in many individuals from several populations, an approach applied in island phylogeography (Kuo et al., 2015; Rodríguez et al., 2013). There are, however, numerous phylogeographic data sets whose sampling is comprehensive, yet sparse. They cover more or less evenly most of the distribution range, but consist of one to a few individuals per population and, at best, a handful of loci. In such cases, no estimates of local genetic diversity are available and conclusions about range expansions are largely based on researcher’s intuition and common wisdom.

## Genetic hubs

Here, I present a method that approximates trends in genetic diversity by integrating information over such globally comprehensive, yet locally sparse sampling. The local diversity is approximated by a distance that *must not* to be travelled from a particular place to access all the observed genetic variation. If all alleles are present at the site, a virtual agent must not to take any travel to access the whole variation. The more depleted is gene pool at the site, the longer are travels of the agent, especially if the site is at the periphery of species distribution. The algorithm is called Genetic hubs as the place of highest diversity (=”genetic hub“) has the same property of “being close to everything” as a hub in public transport network.

In practice, the species distribution is represented by a spatial graph whose vertices correspond to sites and edges to travels between them. The diversity score is estimated separately for each locality. First, it is figured out, which alleles are missing there and what is the shortest path to each of them. Then the graph is effectively reduced just to edges that appear in any of the shortest paths and each edge in the reduced graph is assigned its own weight, which is equal to proportion of variation accessed through the edge. For instance, if the edge appears on the shortest paths to one out of four alleles, its weight is 0.25. The road to be taken for all the remaining genetic variation is then calculated as 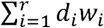, where *d*_*i*_is the length of i^th^ edge, *w*_*i*_ is its weight and the summation goes across *r* edges of the reduced graph. This sum is contrasted to its theoretical maximum, which is calculated in the same way, but assuming all variation at the very most distant locality and no variation at the locality in question. This is equivalent to 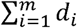 sum of lengths of *m* edges forming the shortest path to the most distant site. Finally, the diversity score is calculated as 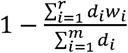. In other words, the length of travel to be taken by the agent is calculated, expressed as a proportion of its theoretical maximum and subtracted from unity to make the value proportional (not inversely proportional) to local diversity. The calculation of diversity score is demonstrated in Figure 1.

**Figure 1.**
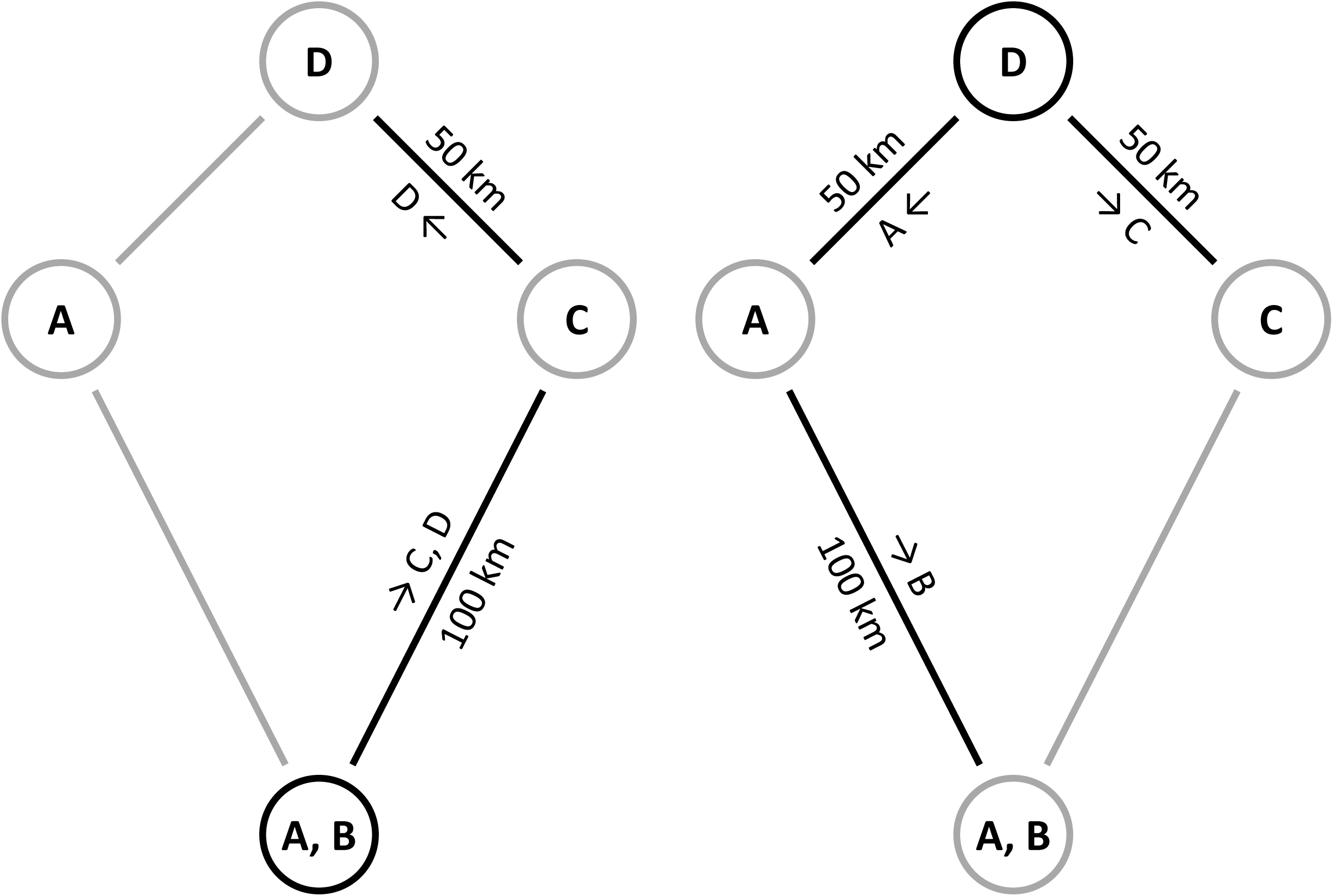
Example of diversity score calculation. Four sites (southern, western, northern and eastern) are shown, each with a different sample of A, B, C, D alleles. In the left panel, the arrows show the shortest paths from the southern site (with A, B alleles) to sites with other alleles. Every allele represent one fourth of total variation and the longer link is on the shortest path to two alleles (C, D), whereas the shorter link on the path to just a single allele (D). The diversity score is therefore calculated as 1 – (100 * 0.50 + 50 * 0.25) / 150 = 0.58. In the right panel, the same is shown for the northern site, whose diversity score turns out to be 0.50.

An obvious, and often essential, extension of the algorithm is to take dissimilarities between alleles into account. This is achieved by modification of the weight calculation, where the proportion of variation accessed through the edge is calculated not only from presences and absences of alleles but also from their dissimilarity matrix. The matrix is subjected to multidimensional scaling and every allele is assigned a vector of values that specify its position in the resulting multidimensional Euclidean space. The space dimensionality is equal to the number of positive eigenvalues in spectral decomposition of the matrix. If the dissimilarity measure is a true metric, all variation can be represented in this space, otherwise some information is lost. In fact, the decision whether or not to weight edges by allele dissimilarities is a delicate one. If we assume recent and fast spread from a single panmictic population, allele dissimilarities do not tell us anything about the expansion. All alleles were well mixed at the beginning and those retained in the same or a nearby population may be very similar as well as largely different from each other. On the other hand, if the average rate of expansion is slow enough to be comparable with effective mutation rate, new alleles are arising along the way and their similarity bears imprint of the expansion process.

Another issue is the choice of spatial graph. The first obvious possibility is the fully wired graph, where all sites are direct neighbors separated just by their physical distance on the Earth surface. It should be appreciated, however, this is not a ‘neutral’, ‘uninformative’ or even ‘universal’ choice. The reason is that any edge passing through inhospitable environment introduces bias as it spuriously indicates contact where none exists. An ideal choice would be probably a fully wired graph with edge lengths modified to reflect landscape resistance to migration (McRae, 2006). Information about the long-term landscape resistance is not easily available, however. A viable alternative is therefore to modify graph structure, either *ad hoc* or according to some predefined criterion, to avoid long-ranging shortcuts that are more risky to introduce substantial bias. The only requirement is the graph has to remain fully connected so that every vertex can be reached, directly or indirectly, from any other vertex. In any case it is advisable to decide about the graph structure prior to any calculation.

The knowledge of genetic hub location is useful especially if it is not contingent on presence of a single sample but supported by the pattern of diversity as a whole. Thus, it is advisable to rerun the algorithm on reduced data sets with sample present at the genetic hub site or at the neighboring sites omitted. This is in fact a jackknifing procedure, although limited to the neighborhood of the genetic hub. Every jackknifed sample is omitted, once at time, but its site is retained as empty one so even after omission of the hub sample itself, the hub’s position may be unchanged. When jackknifing is completed, it is useful to compare genetic hub locations and quantify congruence of diversity trends, e.g. by Spearman correlation coefficient calculated on diversity scores. The omission of samples from sites far apart from the hub is unlikely to change its position and their inclusion into jackknifing could in fact create spurious support for the genetic hub location.

Although the possibility of using various dissimilarities and spatial graphs makes the method remarkably versatile, it has also its inherent limitations. Most importantly, the local genetic diversities are not estimated independently of each other, but instead, they are approximated under the assumption of a single range expansion (with no range expansion as a special case). If the spatial pattern was created, for instance, by population growth at some places and population decline at others, it would not be appropriate to estimate local diversities by integrating information across the whole area. In such case the local diversity is determined by local factors and has to be estimated from local data. Another common case when the algorithm is misled by its crucial assumption is the secondary contact of two expanding populations. Here, the hotspot of diversity is at the frontlines rather than at the origins of the expansions. Due to this limitation Genetic hubs do not allow formal comparison of different historical scenarios. The algorithm also pinpoints just a single site as the hub, although it is likely that ancestral population occupied some larger area. The hub can be thought, therefore, as a centroid of the ancestral range. Finally, and most obviously, more or less even sampling intensity across the whole area is assumed, otherwise the peak of local diversity may be an artefact. The weighting by allele dissimilarity makes the method more robust in this respect.

Genetic hubs are available in the form of open-source package GenHubs for R (R Core Team 2018), which is accessible via CRAN (…). It allows to estimate genetic hub location under a range of settings, namely using different spatial graphs with or without weighting by allele dissimilarity. Apart from the core GenHubs function it offers also jackknifing procedure and associated plotting methods.

## Examples

Overall, Genetic hubs are intended mostly for data exploration and visualization. They are expected to be useful for at least three purposes: (1) a visualization making researcher’s impressions explicit; (2) a preliminary hypothesis formulation where the goal is to pinpoint areas and populations worth of more detailed study; (3) comparative analyses of multiple species or loci where repeatedly spotting the same place as a center of expansion gives higher credit to the implicit assumption of a single expansion scenario. Note also that the virtual agent can travel from places with no data. The algorithm can be therefore used in a predictive manner to estimate local diversities at spots where the species occurrence is known or assumed but from which samples are not available. This feature enhances usefulness of Genetic hubs for the preliminary hypothesis formulation as it allows to assess, which non-sampled locality might be of the greatest interest.

First, we demonstrate usefulness of Genetic hubs on a well-studied example of post-glacial colonization of Europe. Comparative phylogeography of various vertebrate species suggested putative refugia to be located in the Mediterranean (Hewitt, 1999; Schmitt, 2007; Taberlet, Fumagalli, Wust-Saucy & Cosson, 1998), but also in more northerly regions (McDevitt et al., 2012; Kotlík et al., 2006). This view is supported by the fossil record (Knitlová & HoráČek, 2017; Sommer & Nadachowski, 2006; Tougard, Renvoise, Petitjean & Quere, 2008).

Two species of hedgehogs (*Erinaceus*) live in Europe: the Northern White-breasted hedgehog (*E. roumanicus*) lives in the east of Europe, Balkans and in the central Europe, where it meets with its western counterpart, the Western European Hedgehog (*E. europaeus*). In addition there is a pronounced phylogeographic pattern within *E. europaeus* with three distinct mitochondrial lineages (Seddon, Santucci, Reeve & Hewitt, 2001). The whole pattern was interpreted as a result of colonization from refugia located in the Balkans (*E. roumanicus*), Italy (E1 lineage of *E. europaeus*) and Iberia (E2 lineage of *E. europaeus*). The third lineage of *E. europaeus* is confined to Sicilia and won’t be further considered here. The data set reanalyzed here consists of 423 georeferenced records of 100 haplotypes of 426 bp long sequences of mitochondrial control region, originally published by Seddon et al. (2001; 2002), Bolfíková and Hulva (2012) and Černá Bolfíková et al. (2017). The calculation was done in both unweighted and weighted fashion, the latter being based on Kimura two-parameter distances (Kimura, 1980) between haplotypes. In place of spatial graph I used Gabriel graph (Gabriel & Sokal, 1969) with some links crossing the sea manually deleted. Results of the genetic hub analysis are presented in Figure 2. The genetic hub of E2 lineage of *E. europaeus* is located either in Iberia (weighted variant) of in southern France (unweighted variant) as expected from the existing biogeographic scenarios. Location of the genetic hub was surprising in the E1 lineage (in both variants) as it was found in southern Germany instead of Italy. This does not indicate, however, Italy was not a refugium of *E. europaeus* during the last glacial. It only suggests the population from which the rest of the distribution range of E1 lineage was colonized could live more northerly. It is also worth to consider an effect of unbalanced sampling as there are much fewer sites in Italy. The genetic hub of *E. roumanicus* (in both variants) was at the Adriatic coast of Croatia, in accord with the assumed Balkan location of the glacial refugium. In the weighted variant the gradient of diversity is apparent especially in E2 lineage of *E. europaeus* (in the expected north-eastern direction), but in the other two units it also shows interpretable features. In the E1 lineage its minimum is in Scandinavia which is sure to be colonized late and in *E. roumanicus* it has its minimum at the very east suggesting eastward colonization of regions that were inhospitable for long time due to its continental climate. In the unweighted variant (not shown) Iberia also appeared to be colonized late by E2 lineage, which is arguably an artefact caused by not taking allele dissimilarities into account.

**Figure 2.**
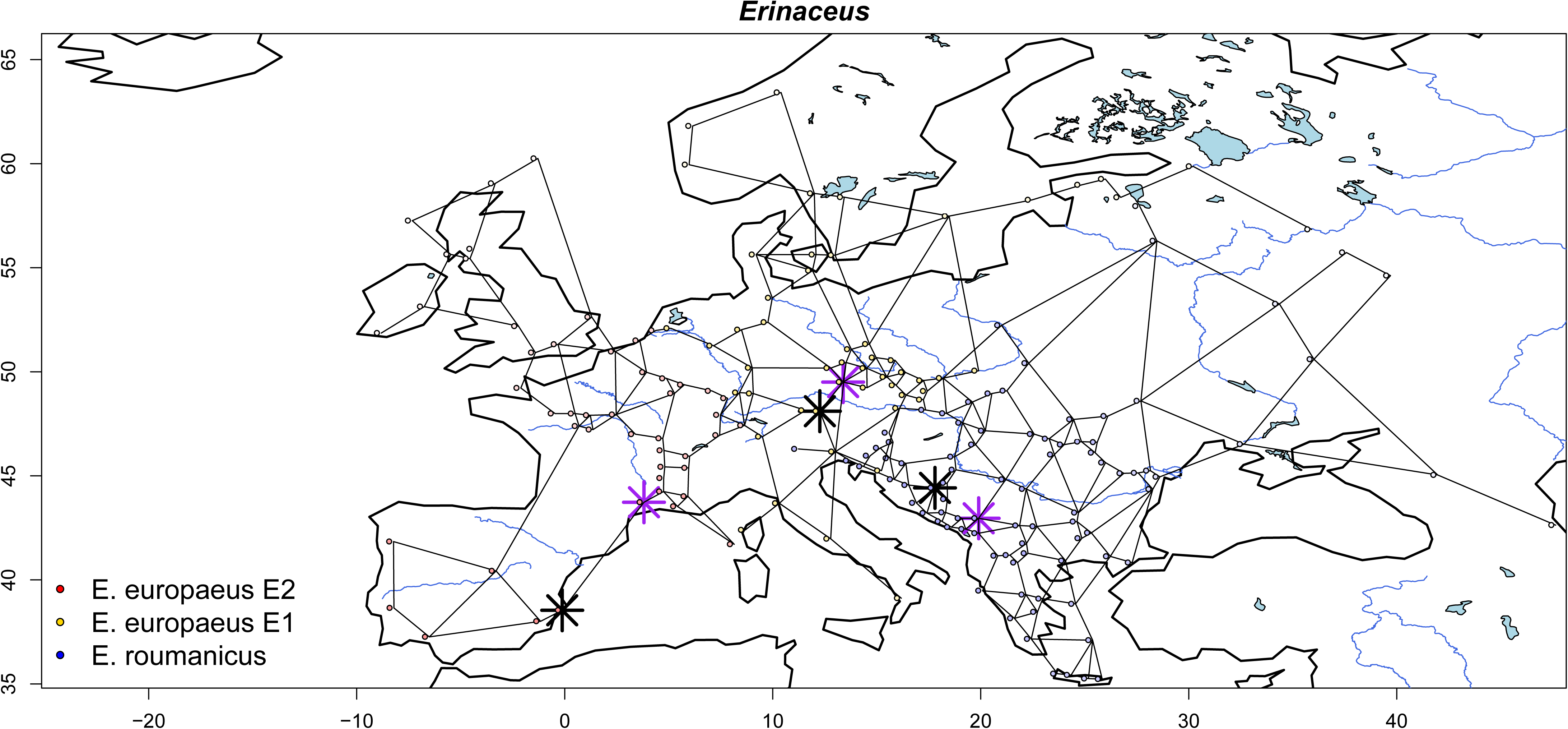
Genetic hubs of three species and lineages of hedgehogs (*Erinaceus*). The shades of colors indicate gradients of diversity for the weighted analysis, whose genetic hub is marked by the black star. Genetic hub from the unweighted analysis is marked by the purple star. Links belong to the three (partially overlapping) graphs upon which the calculation was based.

The phylogeographic structure of the Wood Mouse (*Apodemus sylvaticus*) also likely results from postglacial colonization. Using 981 cytochrome b sequences, Herman et al. (2017) identified six phylogeographic lineages, three of which are analyzed here. The south-eastern lineage is distributed in Italy and Balkans, suggesting glacial refugium somewhere in that regions (Michaux, Magnanou, Paradis, Nieberding & Libois, 2003), the central lineage dominating the western part of continental Europe might spread from a refugium in Iberia or southern France (Michaux et al., 2003) and the newly discovered peripheral lineage is distributed in the British Isles and the eastern Europe which was interpreted to be due to replacement by the central lineage (Herman et al., 2017). Overall, Herman and co-workers were sceptic about possibility of locating glacial refugia from their data set. Indeed, in spite of being impressive in size it comprised only a single mitochondrial locus, which inevitably bears only limited information about population-level processes. However, it is exactly that kind of data set for which Genetic hubs are suited best and where they can bring answer that is provisional, but obtained in a transparent and reproducible manner. The re-analyzed data set includes 445 georeferenced records of 383 cytochrome *b* haplotypes (1140 bp long) from three out of six phylogeographic lineages. The other three were either narrowly localized (Sicilia, Channel Islands) or extraterritorial (northern Africa) and were not considered here. The procedure was the same as in the hedgehog data set. As may be seen in Figure 3, central and south-eastern lineages have their genetic hubs in the presumed refugial regions, southern France and Italy, respectively. In the weighted variant gradient of diversity is well apparent (Figure 3), but the same was the case in unweighted variant (not shown here). Interpretation of the genetic hub location is more complicated in the case of the peripheral lineage. The algorithm unequivocally pinpoints sites in Wales (weighted variant) or even Scotland (unweighted variant), that is in regions which are expected inhospitable for wood mice until the beginning of the Holocene. If the replacement hypothesis of Herman et al. is true, however, the history of these populations is at odds with assumptions of the method, because in such case the geographic pattern of variation was not shaped only by expansion, but also the subsequent wave of local extinctions. Therefore, the genetic hub cannot be interpreted as coincident with the origin of expansion.

**Figure 3.**
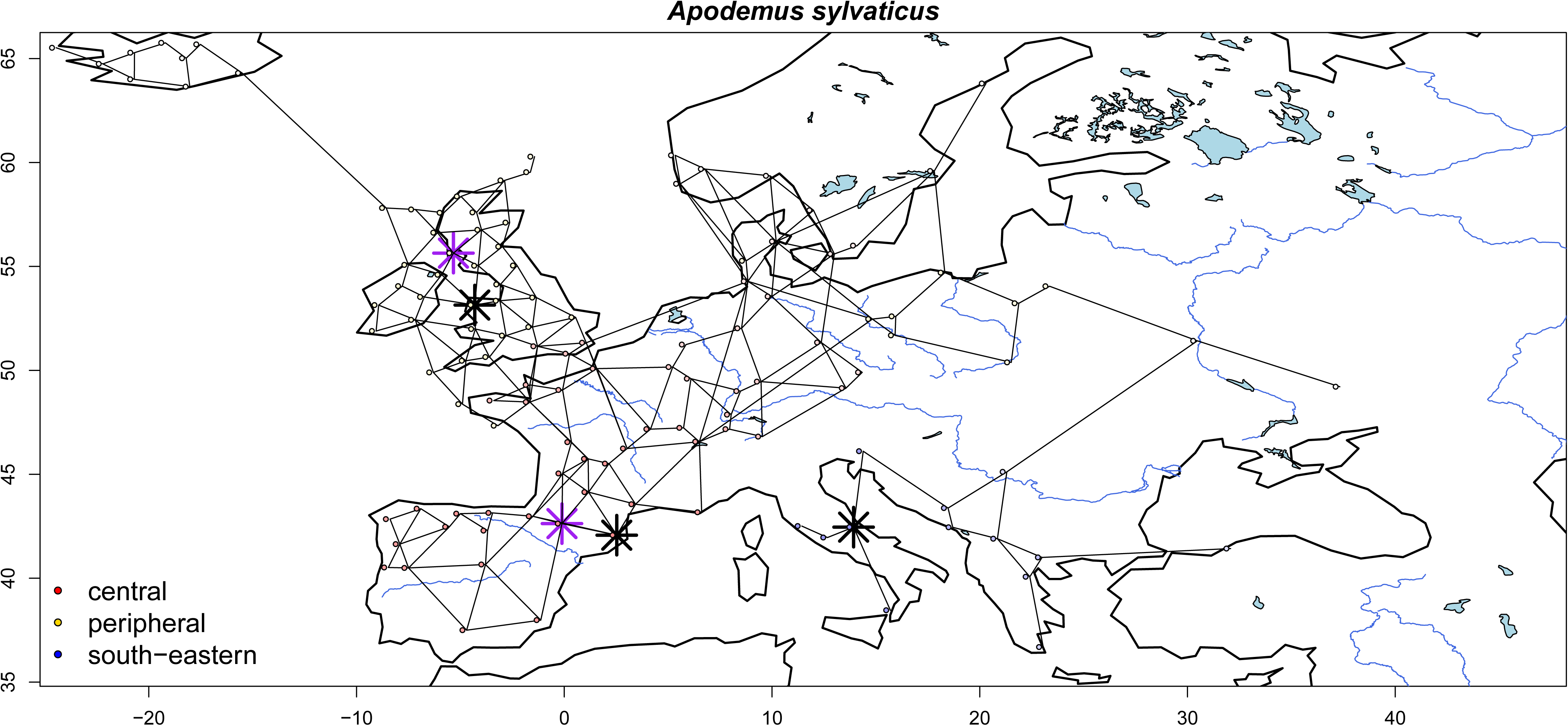
Genetic hubs of three lineages of wood mouse (A. sylvaticus). The sampling sites of the peripheral lineage are not shown completely due to overlap with the other two lineages. Symbols are the same as in Figure 2.

In these two examples Genetic hubs served to examine whether geographical distribution of genetic variation fits *a priori* expectations. More explorative (rather than confirmatory) use of the algorithm is illustrated by a small comparative study, which involves three rodents associated with forests and woodlands in the Tanzanian Eastern Arc Mountains – namely *Grammomys surdaster, Mus triton* and *Praomys delectorum*. The Eastern Arc is a chain of more or less isolated mountain massifs surrounded by savanna, today often turned into agricultural landscape, while in higher elevations they are covered by forests (Platts et al., 2011). More specifically, the focus was on the southern part of the chain as the species considered are absent or represented by genetically distinct lineages in more northerly massifs.

The analyzed data sets were relatively small (18–36 georeferenced records of 16–30 cytochrome *b* haplotypes), but they still allow explicit phylogeographic hypotheses to be formulated (Figure 4). I used the same options as before, except for the Gabriel graph was not manually modified and only weighted genetic hubs were calculated. The data were taken from several published studies (Bryja et al., 2014; 2017; Krásová et al., 2018; Sabuni, Aghová, Bryjová, Šumbera & Bryja, 2018) and supplemented by new records (sequences available in GenBank, accession numbers: XXXXXXXX– XXXXXXXX). Genetic hubs of *G. surdaster* and *P. delectorum* are in the north, while genetic hub of *M. triton* is in the south of the mountain chain. Note, however, that distribution of variability in *M. triton* is not entirely monotonic which calls into question the assumption of a single range expansion as the only force operating behind. In contrast, patterns observed in the other two species seem to be pronounced and monotonic, which is consistent with relatively recent expansion in the southwards direction.

**Figure 4.**
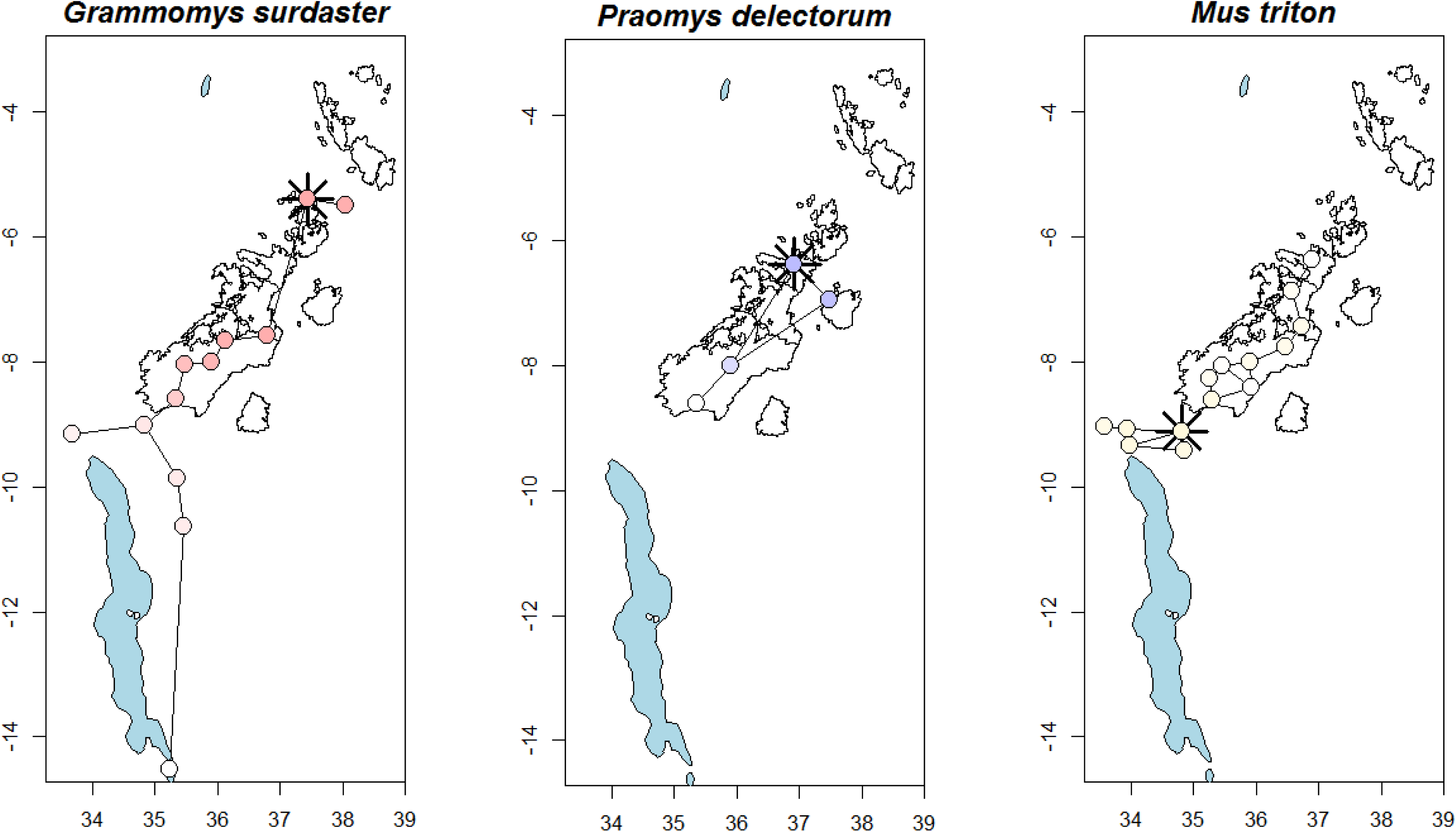
Genetic hubs of three rodent species living the forests and woodlands of the Eastern Arc Mountains.

Jackknifing largely supported the location of genetic hubs in the cases of *Erinaceus* and *A. sylvaticus*. It was conducted with the second order neighborhood, i.e. with sites that were in the graph either directly linked to the genetic hub or separated by at most one other site. In *Erinaceus*, only a few of the jackknifed genetic hubs were displaced from their original location and they always stayed in the same region (not shown). Also the rank correlation between original and jackknifed scores was always very high (ρ≥0.99). In *A. sylvaticus* the correlation was also high: ρ≥0.98 in the central and peripheral lineages and ρ≥0.84 in the south-eastern lineage. Again, only a minor proportion of jackknifed genetic hubs were shifted in location and only one of them deserves special attention. Namely, in both weighted and unweighted variant of the south-eastern lineage analysis, one out of six jackknife replicates resulted in the shift of genetic hub across the sea, from central Italy to the coast of Montenegro.

Results calculated for the three Eastern Arc Mountain species were less robust (Figure 5). The jackknifing here was conducted with just the first order neighborhood (=direct neighbors in the graph), but it still resulted in substantial shifts of genetic hub location. In both *G. surdaster* and *P. delectorum* two out of three jackknife replicates were substantially shifted southward and in *M. triton* two out of four were shifted northward. Score correlations might be as low as 0.50 in *P. delectorum* and 0.53 in *M. triton*, although they were reasonably high (ρ≥0.90) in *G. surdaster*. The results are therefore more dependent on the particular samples in hand and more individuals as well as sampling sites should be employed to obtain more reliable estimates of genetic diversity gradients.

**Figure 5.**
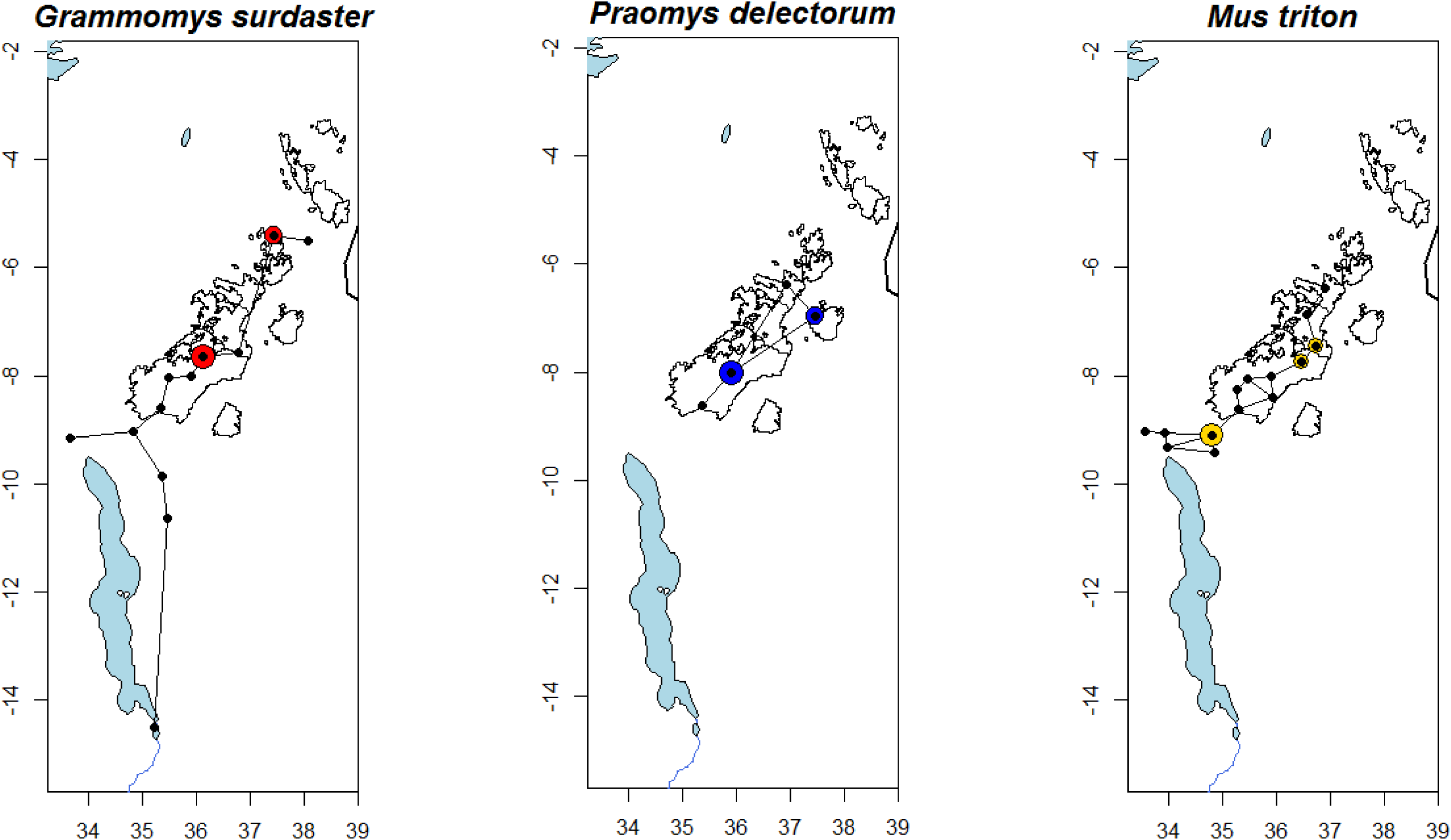
Jackknifing of genetic hubs of the Eastern Arc species. The size of the colored circle indicate the proportion of jackknifed genetic hubs estimated to lie at the particular site.

## Discussion

Genetic hubs method is introduced here as a tool for exploratory data analysis for phylogeography, but also landscape genetics. The two disciplines have similar goals, but they work on different time scales. Landscape genetics is focused on the present and very recent past and thus it assumes spatial arrangement of populations to be more or less static, ancestral alleles to be possibly present in data and spatial variation to be determined mostly by segregation and ongoing migration. If the signal of expansion is present in such data, its origin can be approximated by the unweighted variant of the genetic hub analysis, i.e. without taking allele dissimilarities into account. Phylogeography, instead, is focused on historical processes. It assumes populations shifted largely in location on the time scale of interest, ancestral alleles to be already replaced by their mutational variants and spatial variation to be strongly affected by colonization-extinction process. In such case, the weighted variant of Genetic hubs is more appropriate.

As already mentioned, the most powerful techniques for inference of colonization routes require either multiple individuals to be sampled from every population or a large number of genomic markers to be sampled from every individual. If there was a large set of populations with precise estimates of genetic diversity, one could use any spatial interpolation method (e.g. Miller & Wood, 2014) to identify gradient of variation and its peak. This is seldom the case, however. Alternatively, the origin of range expansion can be also identified from asymmetries in spatial distribution of binary alleles (Peter & Slatkin, 2013), which requires large number of markers but not so many individuals per site. If there are few populations with multiple individuals structured coalescent (aka isolation-with-migration) models (Beerli & Felsenstein, 2001; De Maio, Wu, O’Reilly & Wilson, 2015; Kühnert, Stadler, Vaughan & Drummond, 2016) can be used to identify the source population. If there are few individuals per population, but with a large number of loci genotyped, one can use admixture modelling to get an idea which population was the ancestral one. This can be achieved by proper interpretation of clusters delimited on the basis of Hardy-Weinberg expectations (Pritchard, Stephens & Donnelly, 2000; Guillot, Estoup, Mortier & Cosson, 2005) or by exploiting information about physical location of loci on chromosomes and analysis of introgression blocks (Hellenthal et al., 2014).

Genetic hubs may be used for exploration of any data, which can be converted to lists of alleles present at particular sites (unweighted variant) or to distances between unique genotypes and ultimately components of variation attributable to them (weighted variant). Nevertheless, the method is arguably most useful when dealing with data sets that cannot be analyzed by the abovementioned methods. This is the case for mitochondrial phylogeography that flourished for two decades since 1990 and still represents an important initial step in biogeographical and systematic studies. Typical sampling here is globally comprehensive, but locally sparse: it often includes tens of sampling sites covering substantial part of the species distribution, but only one to a few individuals per site. Whereas the high number of sites precludes the use of parameter-rich structured coalescent, the low number of loci precludes the use of methods that rely on genomic sampling and sequencing of a few more loci does not change the matters substantially. In contrast, genetic hub analysis may provide meaningful results even in such unfavorable situation. Although any single locus carries only partial information about population level processes, some loci can be informative on their own and their conflicting signals may be interpretable as it is the case for maternally, paternally and bi-parentally inherited markers (Toews & Brelsford, 2012).

Nevertheless, there is yet another methodology for ancestral location inference, able to process the very same data sets as just discussed. Lemey, Rambaut, Drummond and Suchard (2009) treated location as a discrete trait, whose evolution unfolds along with the phylogeny itself. In other words, the migration between geographic sites is modelled in the very same way as mutation process. In continuous space an analogous approach treats location as a bivariate trait evolving by diffusion over a plane (Lemey, Rambaut, Welch & Suchard, 2010) or on a sphere (Bouckaert, 2016). Both approaches can be further bridged by modelling migration as a diffusion over a graph of sites covering densely the area suitable for migration (Bouckaert, Bowern & Atkinson, 2018). This methodology, implemented in the Bayesian framework, is open to numerous extensions and modifications including informative priors on particular rates, fixing some of them to zero and allowing them to be asymmetric (Bouckaert et al., 2018; Edwards et al., 2011; Lemey et al., 2009). As such it holds a promise to provide probabilistic estimates of ancestral locations, not only algorithmic ones as provided by Genetic hubs. It remains to be integrated, however, with models taking into account population effective sizes (Kühnert et al., 2016). When population size is omitted, presence of highly divergent haplotypes at the same site is implicitly interpreted as the result of intensive migration rather than as the remnant of ancestral polymorphism. This simplification can greatly bias results, although it may be appropriate for viruses whose mutation and migration rates are comparable (Lemey et al., 2009; Pybus et al., 2012) or for organisms, whose population sizes at remote discrete sites are very small compared to those between the sites (e.g. polar bears analyzed by Edwards et al., 2011).

In summary, the probabilistic inference of ancestral populations and colonization routes is possible when population and/or genomic sampling is intensive and it might become possible for data sets with extensive sampling as well. Genetic hubs method focuses on a part of this problem, namely, on the location of the expansion origin. In this respect, it wants to serve the same purpose in phylogeography as the neighbor-joining method (Saitou & Nei, 1987) does in phylogenetics. It is intended as an approximate, yet fast and versatile, alternative to model-based methods, useful especially for exploratory and preliminary analyses, which can provide good starting points for future studies.

## Acknowledgements

Financial support for this study was provided by the Czech Science Foundation, project no. 18-17398S. I thank to J. Bryja for commenting the first version of the manuscript.

